# Kinesin-14 HSET and KlpA are non-processive microtubule motors with load-dependent power strokes

**DOI:** 10.1101/2023.06.09.544415

**Authors:** Xinglei Liu, Lu Rao, Weihing Qiu, Arne Gennerich

## Abstract

Accurate chromosome segregation during cell division relies on coordinated actions of microtubule (MT)-based motor proteins in the mitotic spindle. Kinesin-14 motors play vital roles in spindle assembly and maintenance by crosslinking antiparallel MTs at the spindle midzone and anchoring spindle MTs’ minus ends at the poles. We investigate the force generation and motility of the Kinesin-14 motors HSET and KlpA, revealing that both motors function as non-processive motors under load, producing single power strokes per MT encounter. Each homodimeric motor generates forces of ∼0.5 pN, but when assembled in teams, they cooperate to generate forces of 1 pN or more. Importantly, cooperative activity among multiple motors leads to increased MT-sliding velocities. Our findings deepen our understanding of the structure-function relationship of Kinesin-14 motors and underscore the significance of cooperative behavior in their cellular functions.

## Introduction

Accurate chromosome partitioning during cell division relies on the proper assembly and maintenance of the mitotic spindle, which is composed of polar and dynamic microtubule (MT) polymers^1^. MT-associated motors such as cytoplasmic dynein, mitotic kinesins, and non-motor MT-associated proteins (MAPs) play crucial roles in the segregation of sister chromatids toward opposite spindle poles^2,3^. Cytoplasmic dynein interacts with astral MTs at the cell membrane and separates duplicated centrosomes at the beginning of mitosis with the assistance of Kinesin-5 (Eg5 in humans and BimC in *Aspergillus nidulans*), which pushes the interacting interpolar MTs of opposite polarity apart. To counterbalance the outward directed forces generated by Kinesin-5 and prevent excessive separation of the two opposing MT arrays while the spindle forms, Kinesin-14 motors (HEST in humans and KlpA in *A. nidulans*) generate the opposing inward directed forces^2,4,5^.

Unlike the processive movement of some members of the Kinesin-5 family towards the plus-end of MTs^6,7^, Kinesin-14 family members are non-processive, minus-end-directed motors. In addition to their C-terminal motor domains, Kinesin-14 motors possess an N-terminal MT-binding tail crucial for their MT cross-linking and sliding function^8–10^. These motors have diverse roles in the assembly and regulation of the mitotic spindle. For example, *Schizosaccharomyces pombe* Kinesin-14 Pkl1 localizes to spindle poles to anchor the minus ends of spindle MTs, while *S. pombe* Kinesin-14 Klp2 crosslinks antiparallel MTs from opposite poles within the midzone to counteract the outward pushing forces generated by Kinesin-5 Cut7^8^. HSET and *Xenopus laevis* Kinesin-14 XCTK2 affect spindle length and morphology through MT sliding^11^. Not surprisingly, dysfunction or dysregulation of Kinesin-14 motors can lead to impaired cell division, including defective spindle assemble and erroneous chromosome segregation^12–16^.

Structure-function studies have provided valuable insights into the molecular functions of Kinesin-14 motors^17–26^. However, despite extensive efforts to understand their mechanisms, a consensus regarding the precise molecular function of Kinesin-14 motors remains elusive. For instance, studies have shown that single Ncd molecules are non-processive, minus-end-directed motors, performing a single ATP-triggered power stroke per MT encounter^18,22, 27–30^. On the other hand, KlpA exhibits processive movement towards the MT plus end at the single-molecule level but displays canonical minus-end-directed motility in multi-motor-based MT-gliding and -sliding assays^17^. Furthermore, while single Ncd molecules generate forces of less than ∼0.5 pN^19,23,24^, single HSET molecules have been suggested to generate forces of up to 1.1 pN by taking multiple processive steps along MTs^25^. Consequently, it remains uncertain whether these reported differences are species-specific or arise from variations in experimental conditions such as the choice of assay, buffer conditions, and protein preparations.

Here, by combining mutagenesis studies with single-molecule fluorescence and optical tweezers assays, we demonstrate that full-length HSET and KlpA behave as non-processive motors under load, producing single power strokes per MT encounter. Our findings reveal that HSET generates a power stroke of ∼9 nm under low-load conditions. However, as the applied load increases, the power stroke is hindered, resulting in a reduced displacement of ∼2 nm under an applied load of ∼0.6 pN. This load-dependency of the lever arm, combined with its limited length, provides an explanation for why Kinesin-14 motors generate only ∼0.5 pN of force at the single-molecule level.

Contrary to previous studies indicating a reduction in MT-gliding velocity with an increasing number of motors^20,31^, we demonstrate that multiple HSET/KlpA molecules work synergistically to generate faster velocities in MT-gliding assays. Interestingly, when assembled in oligomers or other multi-motor configurations, these motors exhibit processive motion under both unloaded (our data and refs^17,19,21,32^) and loaded conditions toward the MT minus end and generate forces of 1 pN and more. Consequently, while Kinesin-14 motors act individually as non-processive and weak motors under load, they exhibit processive motion and generate significant forces when functioning in teams.

## Results

### KlpA is non-processive under load

The force-generation capability of the *D. melanogaster* Kinesin-14 Ncd has been extensively studied at the single-molecule level^19,23,24,33,34^. However, the force generation of the *A. nidulans* Kinesin-14 KlpA remains unexplored. Unlike Ncd, which is non-processive and generates a single ATP-triggered working stroke towards the MT minus end per MT encounter^18,22, 27–30^, KlpA is a processive motor that moves towards the MT plus end^17^. KlpA’s processivity critically depends on its N-terminal MT-binding tail, as KlpA lacking the N-terminal tail behaves like a non-processive motor^17^. To investigate KlpA’s behavior under load, we created GFP-KlpA, a full-length KlpA construct with an N-terminal GFP for the coupling of the N-terminal tail to anti-GFP antibody-coated polystyrene trapping beads (Fig. 1A&B and Suppl. Fig. 1A). To assess the force-generation capability of single KlpA molecules, we conducted optical-trapping experiments at motor concentrations where less than 30% of the beads interacted with the coverslip-attached MTs^35^ (Fig. 1B&C). When the tail of KlpA was bound to a 1-μm trapping bead, it generated a force of approximately 0.5 pN towards the MT minus-end (Fig. 1C&D). This force is comparable to, albeit slightly higher than, the force produced by Ncd (where two Ncd molecules linked through a DNA scaffold generate approximately 0.5 pN^19^). Hence, while KlpA’s MT-binding tail domain facilitates processive motility in the absence of load^17^, KlpA behaves as a non-processive motor under load, capable of generating only single power strokes (Fig. 1C, right).

**Fig. 1.**
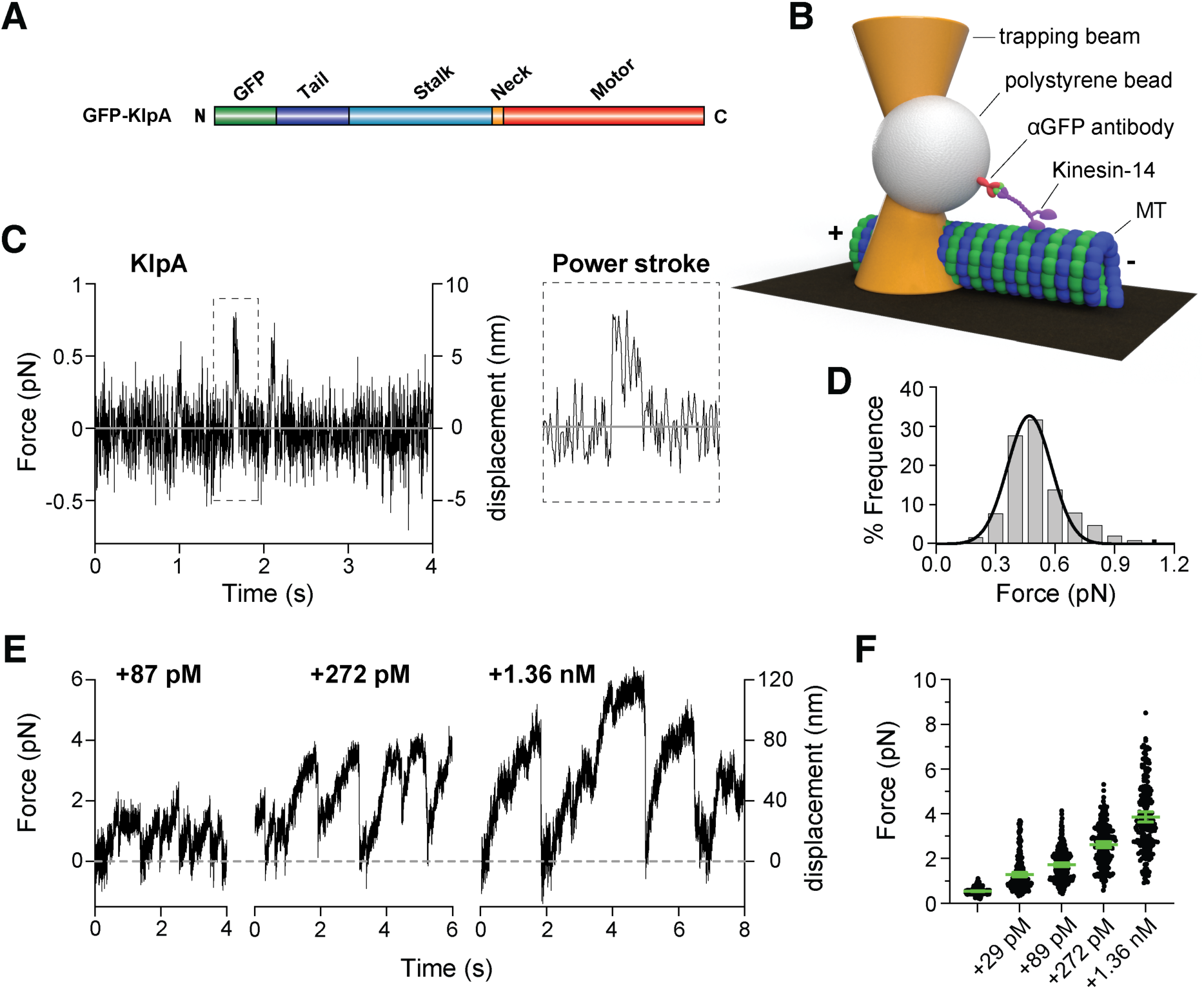
KlpA force generation in isolation and in teams. (A) Structural organization of the full-length recombinant GFP-KlpA construct, highlighting the tail (a.a. 1-152), central stalk (a.a. 153-416), neck (a.a. 417-421), and catalytic motor domain (a.a. 422-756). (B) Schematic representation of the optical-trapping assay, illustrating a polystyrene bead with a bound Kinesin-14 motor trapped by a near-infrared optical trapping beam and positioned above a surface-bound MT (not to scale). (C) Left: Representative force-versus-time trace depicting bead movement driven by a single KlpA molecule at 1 mM ATP and *k*=0.1 pN/nm. Right: Highlighted trace section demonstrating a single-step displacement (“power stroke”) corresponding to the rectangular box on the left. (D) Histogram of forces generated by single KlpA molecules (0.47 ± 0.01 pN, mean ± SEM from Gaussian fit, *n*=493). (E) Representative force traces of GFP-KlpA bound to anti-GFP antibody-coated trapping beads and oligomerized with increasing amounts of added mCherry-KlpA: +87 pM (left), +272 pM (middle) and +1.36 nM (right) mCherry-KlpA. (F) Forces generated by oligomerized KlpA motors at different concentrations of added mCherry-KlpA and *k*=0.05 pN/nm. Green bars represent the mean with 95% confidence intervals (CIs). GFP-KlpA at the single-molecule level (14 pM) without added mCherry-KlpA (see part *D*): 0.51 ± 0.01 pN (mean ± SEM, *n* = 493); +29 pM mCherry-KlpA: 1.29 ± 0.05 pN (*n*=232); +87 pM mCherry-KlpA: 1.73 ± 0.04 pN (*n*=259); +272 pM mCherry-KlpA: 2.62 ± 0.07 pN (*n*=203); and +1.36 nM mCherry-KlpA: 3.86 ± 0.12 pN (*n*=187). Note: a.a. refers to amino acid.

### Multiple KlpA molecules work cooperatively to generates larger forces

KlpA exhibits the ability to form oligomeric complexes consisting of multiple motors that move processively towards the MT minus end^17^. To investigate the cooperative behavior of KlpA under load, we conducted force measurements while varying the concentration of KlpA (while keeping the bead concentration constant). To differentiate between the effects of KlpA oligomerization and the recruitment of individual KlpA molecules to distinct positions on the bead through surface-bound anti-GFP antibodies, we introduced increasing concentrations of mCherry-tagged motors (Suppl. Fig. 1B) to beads pre-coated with GFP-tagged motors. The concentration of GFP-tagged motors was carefully chosen to ensure that less than 30% of the pre-coated beads exhibited force generation events, indicating the presence of only one GFP-tagged motor per force-generating bead^37^. In contrast to Kinesin-1, which shows minimal dependence on the number of motors^19,36^, multiple KlpA motors cooperate to generate forces of 1 pN and higher (Fig. 1E). This cooperative behavior observed in KlpA is reminiscent of Ncd, which also generates additive forces when multiple motors are engaged^19^. Although some non-specific motor-bead interactions were observed at higher concentrations of added mCherry-KlpA (as mCherry does not bind to the anti-GFP antibodies) (Suppl. Fig. 2), our results unequivocally demonstrate the contribution of KlpA oligomerization to the observed increase in force generation upon the addition of mCherry-KlpA (Fig. 1F).

### HSET is a non-processive motor that cooperates when working in teams

In contrast to previous studies on Ncd^19,23,24,34^ and our findings on KlpA (Fig. 1C&D), recent research suggested that HSET, the human Kinesin-14 orthologue, exhibits weak processivity, capable of generating several 8-nm steps under loads up to ∼1.1 pN^25^. To investigate potential differences in behavior under load between KlpA and HSET, we expressed full-length HSET fused with an N-terminal GFP tag (Fig. 2A and Suppl. Fig. 1C) and conducted optical-trapping experiments to examine its force generation. Contrary to the previously reported behavior of HSET^25^ but consistent with our KlpA results (Fig. 1C), we discovered that HSET generates a maximum force of approximately 0.5 pN (Fig. 2C&D). To confirm that these forces originate from single HSET motors, we conducted photobleaching experiments on HSET motors bound to coverslip-attached MTs in a strong MT-binding state (induced by AMP-PNP^18^) (Fig. 2G). The experiments revealed predominantly two photo-bleaching steps, indicating that HSET primarily formed homodimers in solution. A minor contribution of 3 and 4 photo-bleaching steps was also observed, suggesting the possible existence of a small population of HSET tetramers. This observation aligns with our findings for KlpA (shown in Figure 2I) and is supported by our fluorescence microscopy measurements, which showed that HSET formed higher-order particles at higher motor concentrations and exhibited processive movement towards the MT minus end (Fig. 2H and Suppl. Fig. 3A). Alternatively, the observation of 4 photo-bleaching events could be explained by the rare occurrence of two dimeric motors landing within the same diffraction-limited spot on the MT. Subsequently, we measured the fraction of force-generating beads as a function of increasing HSET concentrations while maintaining a constant bead concentration. The resulting curve fit best to a model derived for one or more motors, rather than a model for two or more motors^35^ (Fig. 2B), confirming that a single HSET motor is capable of generating the observed force of ∼0.5 pN. Further examination of the measured force-generation events revealed the presence of single, non-processive “power strokes” (Fig. 2C, right), indicating that HSET, similar to its orthologues KlpA (Fig. 1C, right) and Ncd^23,24^, behaves as a non-processive motor under load.

**Fig. 2.**
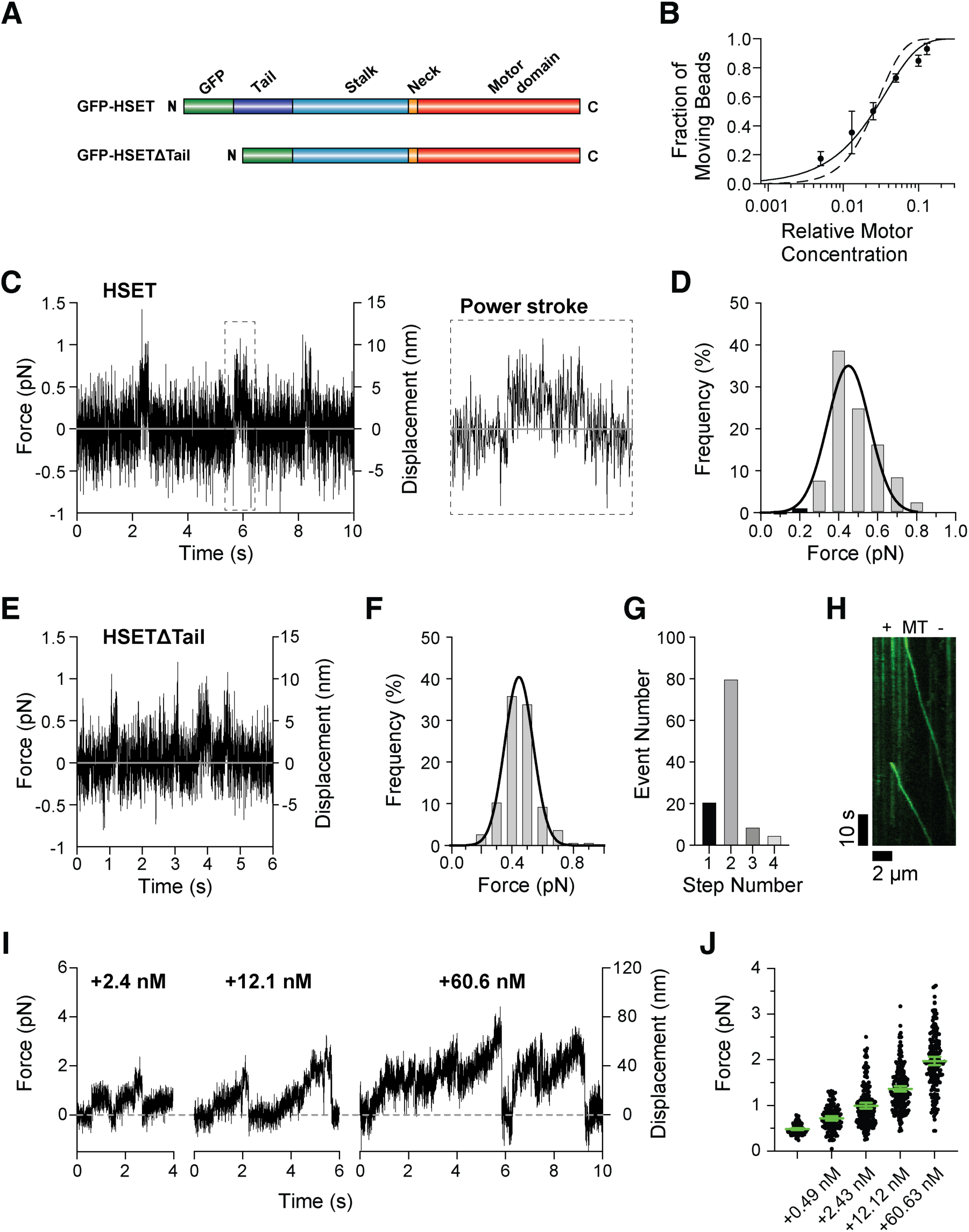
HSET and HSETΔTail behavior under load. (A) Structural organization of full-length HSET (GFP-HSET) and tail-truncated HSET (GFP-HSETΔTail), highlighting the tail (a.a. 1-138), central stalk (a.a. 139-305), neck (a.a. 306-310), and catalytic motor domain (a.a. 311-673). (B) Fraction of GFP-HSET-coated beads binding to and moving along MTs as a function of relative motor concentration. The bead concentration was constant, while the motor concentration was varied (*n* = 321 total number of beads tested; *n* = 30–70 at each concentration). The solid line represents the fit to the Poisson distribution 1 − exp(−λC) for one or more motor molecules, where C is the relative motor concentration and λ is a fit parameter (R^2^ = 0.933). The dotted line represents the fit to the distribution 1 − exp(−λC) − (λC)exp(−λC) for two or more molecules (R^2^ = 0.802). (C) Representative force-versus-time trace of bead movement driven by a single HSET molecule at 1 mM ATP and k=0.1 pN/nm. (D) Histogram showing forces generated by single HSET molecules (0.45 ± 0.01 pN, mean ± SEM from Gaussian fit, *n*=116). (E) Representative force-versus-time trace of bead movement driven by a single HSET-ΔTail molecule at 1 mM ATP and k=0.1 pN/nm. (F) Histogram of forces generated by single HSET-ΔTail molecules (0.45 ± 0.01 pN, mean ± SEM from Gaussian fit, *n*=197). (G) Photobleaching analysis of full-length HSET, showing the number of bleaching events with one (21), two (80), three (9) and four (5) photobleaching steps. (H) Example of MT minus-end-directed processive motion of HSET particles formed at 1 nM motor concentration. These particles consist of multiple HSET motors working together. (I) Representative force traces of assemblies of GFP-HSET molecules at increasing concentrations of added GFP-HSET motors: +2.4 nM, +12.1 nM and +60.6 nM. (J) Forces generated by GFP-HSET motors at varying concentrations of added motors at *k*=0.1 pN/nm. Green bars represent the mean values with 95% CIs. GFP-HSET at the single-molecule level: 0.48 ± 0.01 pN (mean ± SEM, *n*=116); +0.49 nM: 0.72 ± 0.02 pN (*n*=128); +2.43 nM: 1.0 ± 0.03 pN (*n*=201); +12.12 nM: 1.36 ± 0.03 pN (*n*=232); and +60.63 nM: 1.97 ± 0.05 pN (*n*=191).

Based on our observations of single- and multi-motor behavior of KlpA, as well as the finding that single HSET molecules generate only ∼0.5 pN of force, we hypothesized that the reported ∼1.1 pN force of HSET is a result of cooperative activity among multiple HSET molecules. To test this hypothesis, we measured the force generation of HSET at varying HSET concentrations. Indeed, we observed forces of 1 pN and above at higher HSET concentrations (Fig. 2I&J), providing support for our hypothesis that the reported 1.1 pN force of HSET arises from cooperative behavior among multiple HSET molecules. In conclusion, while individual HSET molecules exhibit non-processive and weak force generation behavior under load, multiple HSET molecules work cooperatively to generate larger forces, similar to the behavior observed for multiple KlpA (Fig. 1E&F) and multiple Ncd molecules^19^.

### Full-length and tail-truncated HSET behave indistinguishably under load

A previous study also indicated that the N-terminal MT-binding tail of HSET may affect the motility of HSET-coated beads under unloaded conditions^25^. However, in the optical-trapping assay, where the tail is pulled away from the MT surface due to its coupling to the trapping bead, its interaction with MTs is unlikely. Based on this understanding, we hypothesized that the force generation of HSET without the N-terminal tail would be similar to full-length HSET. To test this hypothesis, we expressed a tail-truncated HSET construct with an N-terminal GFP (GFP-HSETΔTail, Fig. 2A) and examined its force generation. Like full-length HSET, single tail-truncated HSET molecules generated forces of ∼0.5 pN (Fig. 2E&F). Additionally, at the multi-motor level, GFP-HSETΔTail exhibited cooperative behavior to generate larger forces (Suppl. Fig. 3B). These findings demonstrate that both KlpA and HSET exhibit comparable force generation capabilities at both the single- and multi-motor level.

### HSET performs a single power stroke to generate 0.5 pN

The force generated by single Kinesin-14 molecules, such as HSET and KlpA, is significantly weaker (∼0.5 pN; Fig. 1C&D and Fig. 2C&D) than the force generated by Kinesin-1 (∼5 pN^37,38^) and Kinesin-3 (∼3 pN^37,39^). To investigate why HSET and KlpA produce only ∼0.5 pN, we examined the cryoEM structure of Ncd, which suggested that despite having two motor domains, Ncd generates a single ∼9 nm power stroke per interaction with the MT (MT binding, power stroke, followed by detachment) (Fig. 3A). Based on these results, we hypothesized that if the power stroke size remains constant regardless of the applied load, the force output in the trapping assay should increase with higher trap stiffness (corresponding to a higher force per unit displacement). To test this, we measured the force generation of a single HSET molecule as a function of trap stiffness and discovered that the force output indeed increased with increasing trap stiffness up to 0.05 pN/nm (Fig. 3B). However, beyond 0.05 pN/nm, the force generated showed only a moderate increase. This biphasic behavior can be explained by the measured displacements: for trap stiffnesses up to 0.05 pN/nm, the displacement (i.e., power stroke size) was approximately 9 nm, consistent with the cryoEM study on Ncd, and independent of trap stiffness (Fig. 3C, left). The measured force corresponding to a 9 nm displacement at 0.05 pN/nm trap stiffness was ∼0.5 pN, aligning with our experimental measurements (Fig. 2C&D). Thus, the force generated by HSET is primarily determined by the size of its power stroke for trap stiffnesses up to 0.05 pN/nm. However, as the trap stiffness increases further, the movement of HSET’s “lever arm” becomes increasingly restricted, resulting in smaller effective displacements. This is evident from the decreasing displacement per MT encounter, reaching 1.8 nm at 0.3 pN/nm (Fig. 3C&D). Therefore, the full power stroke can only occur under unloaded or low-load conditions (Fig. 3C, left), indicating that the largest displacements achievable by HSET and KlpA can be observed under high loads *in vivo* when multiple motors acting on the same pair of MTs share the generated load.

**Fig. 3.**
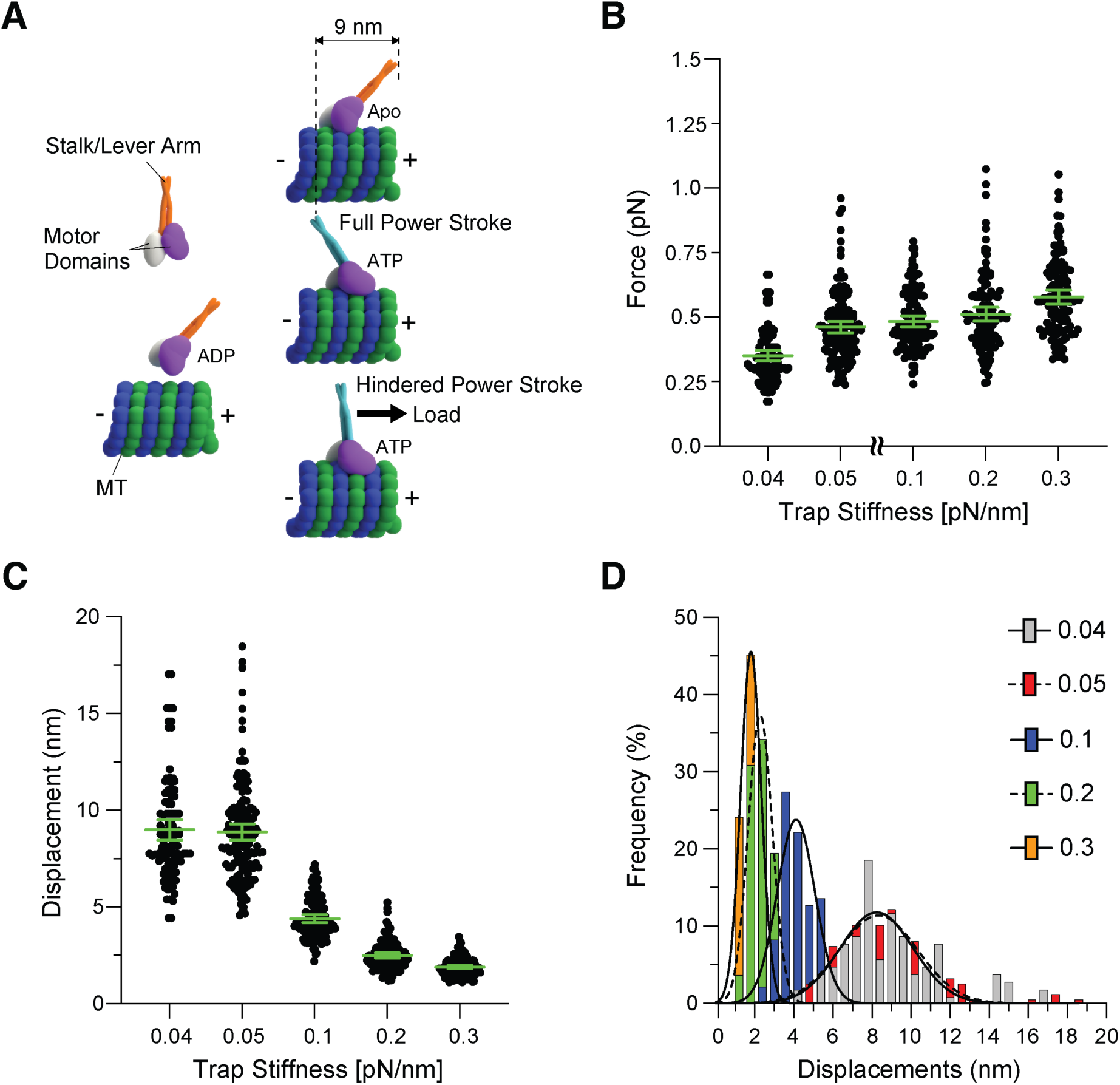
HSET power stroke and force generation. (A) Schematic representation of the ∼9 nm power stroke by HSET under no load or low load conditions, and restrained power stroke under increased loads. (B) Force generated by single HEST molecules as a function of increasing trap stiffnesses. Green bars represent the mean values with 95% CIs. 0.35 ± 0.01 pN (mean ± SEM) at *k*=0.04 pN/nm (*n*=101); 0.46 ± 0.01 pN at *k*=0.05 pN/nm (*n*=145); 0.48 ± 0.01 pN/nm at *k*=0.1 pN/nm (*n*=116); 0.51 ± 0.01 pN at *k*=0.2 pN/nm (*n*=122); and 0.58 ± 0.01 pN at *k*=0.3 pN/nm (*n*=119). (C) Displacement as a function of increasing trap stiffnesses. Green bars represent the mean values with 95% CI. 8.99 ± 0.26 nm at *k*=0.04 pN/nm (*n*=101); 8.88 ± 0.21 nm at *k*=0.05 pN/nm (*n*=145); 4.40 ± 0.1 nm at *k*=0.1 pN/nm (*n*=116); 2.49 ± 0.07 nm at *k*=0.2 pN/nm (*n*=122); and 1.89 ± 0.04 nm at *k*=0.3 pN/nm (*n*=119). (E) Histogram of displacements generated by single HSET molecules as a function of increasing trap stiffnesses. 8.24 ± 0.19 nm at *k*=0.04 pN/nm (mean ± SEM from Gaussian fit); 8.33 ± 0.17 nm at *k*=0.05 pN/nm; 4.12 ± 0.090 nm at *k*=0.1 pN/nm; 2.28 ± 0.096 nm at *k*=0.2 pN/nm; and 1.81 ± 0.048 nm at *k*=0.3 pN/nm.

### HSET molecules work cooperatively to increase the velocity during MT Gliding

Kinesin-14 motors play a critical role in sliding anti-parallel MTs within the mitotic spindle^2^. Unlike processive Kinesin-1 motors^19,36^, Kinesin-14 motors exhibit non-processive interactions and demonstrate cooperative force generation, similar to non-processive muscle myosin 2 that drives muscle contraction^40–42^. In a manner reminiscent of non-processive myosin motors in muscle^42,43^, non-processive Kinesin-14 motors, such as HSET, briefly engage with a MT to perform a power stroke before detaching (Fig. 2C, left). Due to their low duty ratio (the fraction of time the motor spends bound to the filament is less than 0.5), one might expect that the velocity of anti-parallel MTs being slid together (inward) by HSET would increase with the number of motors binding stochastically to MTs. However, studies on both HSET and Ncd have indicated that the velocity in MT-gliding assays (Fig. 4A) decreases as the number of motors increases^20,31^. We replicated this unexpected result using full-length HSET (Fig. 4C). While we immobilized GFP-HSET motors on the cover-glass surface using surface-bound anti-GFP antibodies and took precautions to prevent non-specific surface binding using casein, we suspected that some GFP-HSET motors could non-specifically bind to the surface such that their N-terminal tails could act as brakes by binding to the gliding MTs to slow down gliding (Fig. 4A). To verify our hypothesis, we conducted a control experiment in which full-length HSET was non-specifically absorbed onto the coverslip of the microfluidic slide chamber (without surface-bound anti-GFP antibodies; Fig. 4B). Indeed, we observed MT gliding and a reduction in gliding speed with increasing motor concentrations even in the absence of surface-absorbed antibodies (Fig. 4D), confirming that non-specifically interacting motors contribute to the decrease in gliding velocity.

**Fig. 4.**
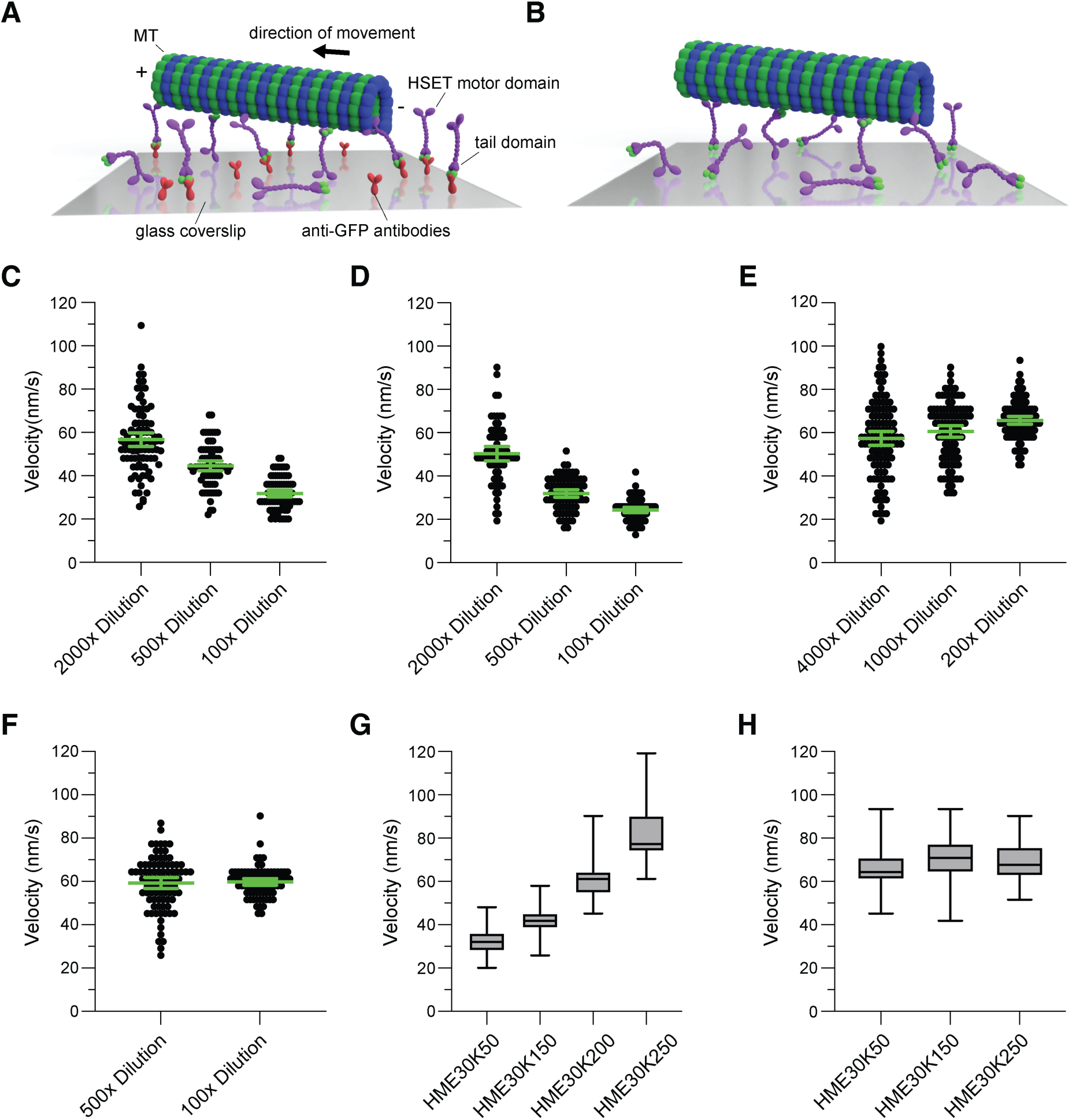
Cooperative effect of multiple HSET molecules on MT gliding velocity. (A) Schematic representation of the MT-gliding assay using anti-GFP antibodies to bind the GFP-HSET to the cover glass. Non-specifically bound motors interact with the gliding MT through their tails. (B) Schematic representation of a MT-gliding assay without surface-bound antibodies, where all motors are absorbed non-specifically to the cover glass. Increased interactions between the motor tails and the gliding MT occur due to random motor orientations. (C) MT-gliding velocities with increasing concentrations (less dilution) of HSET molecules bound to the cover glass via anti-GFP antibodies. Starting concentration: 2.91 μM GFP-HSET. Green bars represent the mean values with 95% CIs. MT-gliding velocity in HME30K50 buffer (HME30 buffer with 50 mM K^+^) at 2,000 dilution: 56.7 ± 1.6 nm/s (*n*=92); 500 dilution: 44.7 ± 1.1 nm/s (*n*=75); and 100 dilution: ± 0.8 nm/s (*n*=81). (D) MT-gliding velocity for non-specifically bound HSET molecules. Starting concentration: 2.91 μM GFP-HSET. Green bars represent the mean values with 95% CIs. MT-gliding velocity in HME30K50 buffer at 2,000 dilution: 50.3 ± 1.7 nm/s (*n*=69); 500 dilution: ± 0.9 nm/s (*n*=78); and 100 dilution: 24.3 ± 0.6 nm/s (*n*=77). (E) MT-gliding velocity for GFP-HSET-ΔTail molecules bound to the cover glass via anti-GFP antibodies. Starting concentration: 4.93 μM GFP-HSETΔTail. Green bars represent the mean values with 95% CIs. MT-gliding velocity in HME30K50 buffer at 4,000 dilution: 57.4 ± 1.6 nm/s (*n*=117); 1,000 dilution: 60.6 ± 1.4 nm/s (*n*=102); and 200 dilution: 65.7 ± 0.9 nm/s (*n*=109). (F) MT-gliding velocity of HSET molecules in HME30 buffer containing 200 mM K^+^. MT-gliding velocity in HME30K200 buffer at 500 dilution: 59.2 ± 1.3 nm/s (*n*=88); and 100 dilution: 59.7 ± 0.8 nm/s (*n*=81). (G) MT-gliding velocity as a function of increasing ionic strength (HME30 buffer with 50 mM, 150 mM, 200 mM or 250 mM K^+^). Velocities are depicted in box and whisker plots. Velocities were measured with 29.1 nM GFP-HSET for each group. MT-gliding velocities measured with HME30K50 buffer: 31.8 ± 0.8 nm/s (*n*=81); with buffer HME30K150: 42.4 ± 0.7 nm/s (*n*=81); with buffer HME30K200: 59.7 ± 0.8 nm/s (*n*=81); and with buffer HME30K250: 81.7 ± 1.4 nm/s (*n*=83). (H) MT-gliding velocities of GFP-HSET-ΔTail molecules bound to the cover glass via anti-GFP antibodies are insensitive to increasing ionic strength. The velocities were measured using 24.7 nM GFP-HSET-ΔTail for each group. MT-gliding velocities measured in HME30K50 buffer: 65.7 ± 0.9 nm/s (*n*=99); with buffer HME30K150: 70 ± 1 nm/s (*n*=102); with buffer HME30K250: 68.8 ± 0.9 nm/s (*n*=97).

Next, we performed a MT-gliding assay using tail-truncated HSET (GFP-HSETΔTail, Fig. 2A) and observed an increase, rather than a decrease, in the velocity of MT gliding with increasing motor concentrations (Fig. 4E), which aligns with the behavior expected for a motor with a low duty ratio. Furthermore, as we increased the ionic strength, which weakens the interaction between the tail and the MT, the effect on MT-gliding velocity became less pronounced at 200 mM KCl (Fig. 4F), and the overall gliding velocity increased with salt concentration (Fig. 4G). Supporting this finding, we observed that the velocity of MT gliding powered by the tail-truncated motor did not increase with increasing ionic strength (Fig. 4H), suggesting that HSET’s tail strongly interacts with MTs through ionic interactions, while the motor domain exhibits weaker interactions and is less sensitive to changes in salt concentration. In the latter case, the motor domain may have a weaker interaction with the MT at higher ionic strength, resulting in increased velocity. However, this increase could be counteracted by a shorter binding time during the power stroke, leading to a decrease in velocity. Thus, multiple HSET molecules work together, with their tails binding to neighboring MTs (rather than the same MT), to enhance both force generation and velocity during MT sliding.

## Discussion

By conducting mutagenesis experiments coupled with single-molecule studies, we have addressed the discrepancies observed in previous studies regarding the force-generation and movement capabilities of Kinesin-14 motors. Our findings demonstrate that both the *Aspergillus nidulans* Kinesin-14 motor KlpA and its human counterpart HSET exhibit non-processive behavior under load, generating only a single power stroke per MT encounter, despite possessing two motor domains. Consequently, the forces generated by individual motor molecules are relatively weak, measuring approximately 0.5 pN. However, when Kinesin-14 motors function collectively as a team, they display remarkable processivity and can generate forces of 1 pN and higher.

The finding that KlpA generates a ∼0.5 pN force towards the MT minus end and behaves non-processively under load (Fig. 1C&D and Fig. 2 C&D) might be unexpected as we previously demonstrated that KlpA moves processively towards the MT plus end in the absence of load^17^. Typically, processive kinesin motors generate forces of 1 pN or more^37,44–46^. However, we have shown that KlpA’s processivity depends on its N-terminal MT-binding tail, as removal of this tail abolishes its ability to move processively^17^. In our trapping assay, the tail of KlpA is bound to a polystyrene bead, mimicking its *in vivo* configuration where the motor cross-links two anti-parallel MTs, with its two motor domains bound to one MT and the tail domains bound to the other MT. Consequently, the tail is unable to interact with the same MT as the motor domains (if the tail would bind to the same MT as the motor domains, the motor would not contribute to MT sliding), explaining the non-processive behavior of KlpA under load.

In contrast to KlpA, both HSET and Ncd are non-processive in single-molecule fluorescence assays^19–21^. However, when a flexible polypeptide is inserted into the central stalk domain of KlpA and Ncd, both motors exhibit processive movement towards the MT minus end^17^. This observation indicates that the ability of these Kinesin-14 motors to move processively not only relies on the MT-binding tails but also on the stiffness or flexibility of the dimerizing stalk^17^. However, once the tail is bound to its cargo, such as a neighboring, antiparallel MT in the mitotic spindle or a polystyrene trapping bead, the stiffness or flexibility of the central stalk becomes irrelevant. Consequently, Kinesin-14 motors behave in a non-processive manner under load.

Moreover, our previous study provided evidence that KlpA has the ability to oligomerize, and these multi-motor complexes exhibit processive movement towards the minus ends of MTs^17^. Consistent with these findings, our force measurements revealed that GFP-KlpA, when bound to a polystyrene trapping bead through an anti-GFP antibody, can oligomerize with mCherry-KlpA upon the addition of increasing concentrations of the latter to the slide chamber. This oligomerization process, as well as the presence of a greater number of non-specifically bound non-interacting motors, both resulted in enhanced force generation towards the minus ends of MTs (Fig. 1E&F). While the processive Kinesin-1 exhibits only a weak dependence on the number of motors^19,36^, KlpA (Fig. 1E&F), HSET (Fig. 2I&J), and Ncd^19^ demonstrate cooperative behavior, leading to increased force output. Based on these findings, we propose that the non-processive behavior of Kinesin-14 motors under load is pivotal for their cooperative ability. Non-processive motors with low duty ratios are less likely to interfere with the activity of other motors compared to processive motors with high duty ratios. Thus, similar to the non-processive myosin 2 responsible for muscle filament sliding^40–42^, Kinesin-14 motors have evolved to work collectively to drive the inward-directed sliding motion of anti-parallel MTs and counteract the outward-directed activity of the mitotic plus-end-directed Kinesin-5 motors, which act on the same pair of MTs, generating movement in the opposite direction to the Kinesin-14 motors^2^.

Previous force measurements on HSET suggested that HSET is weakly processive under load, generating forces of up to 1.1 pN^25^. These forces require HSET to take 7-8 steps under load at the utilized trap stiffnesses. However, through photo-bleaching experiments (to rule out motor aggregation) and motor-dilution experiments using the optical-trapping assay, we conclusively demonstrated that single HSET molecules are non-processive, performing only one power stroke per MT encounter and generating forces of only ∼0.5 pN (Fig. 2C&D). Importantly, we verified that the expression system was not a source of error, as HSET expressed in insect cells and *E. coli* bacteria generated similar ∼0.5 pN forces (Fig. 2C&D and Suppl. Fig. 4). Our findings strongly suggest that the previously reported 1.1 pN force may have resulted from the cooperative activity of multiple HSET molecules. This notion is further supported by the observation that a force of ∼1 pN requires the cooperation of 2-3 Ncd molecules^19^.

At higher motor concentrations, we observed substantial forces up to ∼8 pN for KlpA and ∼4 pN for HSET. The measured forces depend on factors such as the maximum number of motors in an oligomer and the presence of motors that non-specifically bind to the bead surfaces, contributing to force generation. These forces required movements spanning over ∼100 nm under load (Fig. 1E and Fig. 2I). This observation suggests that multiple KlpA and HSET molecules exhibit processive motion when working collectively. The generality of this property within the Kinesin-14 family is supported by studies on Ncd. Ncd not only generates additive forces when functioning in teams but also demonstrates processive movement under unloaded conditions when assembled in groups using DNA scaffolds^19^. Additionally, assemblies of the *Saccharomyces cerevisiae* Kinesin-14 motor Pkl1 exhibit processive motion along MTs when bound to a single quantum dot^32^. Hence, the cooperative behavior observed among non-processive Kinesin-14 motors appears to be conserved across different species.

The limited force production of Kinesin-14 motors can be explained by a cryoEM study that investigated the ATP-driven power stroke of Ncd, revealing a ∼9 nm displacement^18^. In contrast to the processive movement of Kinesin-1, which takes ATP-hydrolysis driven hand-over-hand steps of ∼8 nm along MT filaments^47^, Ncd exhibits a distinct mechanism. During each encounter with a MT, Ncd’s coiled-coil stalk undergoes a single ATP-driven rotation of ∼70° towards the minus-end of the MTs, resulting in a finite swing of the lever arm and a displacement of ∼9 nm along the long MT axis. The force production is limited by both Ncd’s ability to move the lever arm forward under load and the force generated per displacement. Under low load conditions, HSET’s force output increases with increasing trap stiffness, reaching a maximum at 0.05 pN/nm (with an underlying displacement of ∼9 nm). However, at higher trap stiffnesses, the generated power stroke decreases, indicating that the increased load on HSET prevents it from completing a full power stroke. For instance, at 0.3 pN/nm, HSET generates only a ∼2 nm displacement under an applied load of ∼0.6 pN (Fig. 3B-D). Consequently, the limited length and load-dependency of their lever arms result in KlpA and HSET generating a maximum force of ∼0.5 pN. Although it has been suggested that two Ncd molecules are required to achieve ∼0.5 pN force^19^, the utilized low trap stiffnesses (0.014-0.027 pN/nm) likely hindered the observation of the full force-generation capabilities of single Ncd molecules. In fact, at 0.04 pN/nm, we already observed a reduced force of 0.35 pN for HSET (Fig. 3B). In summary, Kinesin-14 motors are non-processive and weak when acting individually under load, but they exhibit processive motion and generate significant forces when working in teams.

Finally, we have successfully identified the underlying cause of the previously observed negative cooperativity between HSET and Ncd multi-motor assemblies in MT-gliding assays^20,31^. We discovered that this phenomenon arises from undesired interactions between the N-terminal MT-binding tails of the Kinesin-14 motors and the transported MT, facilitated by non-specific interactions of the motor with the cover glass. Essentially, these interactions act as brakes, impeding the gliding motion. To validate this hypothesis, we conducted MT-gliding assays using HSET lacking the N-terminal tail. As anticipated, we observed a significant increase in velocity for the non-processive HSET motor as the motor concentrations were increased. Thus, in a manner similar to myosin 2 motors functioning in muscle^40,41^, Kinesin-14 motors have evolved to collaborate effectively when sliding anti-parallel MTs together (inward) within the mitotic spindle. This cooperative behavior generates substantial forces that counterbalance the opposing forces generated by the MT plus-end-directed tetrameric Kinesin-5 motors.

## Acknowledgments

X. Liu was supported by National Institutes of Health grants R01GM098469 and R01GM127922. L. Rao and A. Gennerich were supported by National Institutes of Health grants R01GM098469 and R01NS114636. W. Qiu was supported by National Institutes of Health grant 1R01GM127922.

## Author Contributions

X. Liu produced and purified all proteins and performed the research; X. Liu, L. Rao and A. Gennerich designed research; X. Liu and A. Gennerich analyzed the experimental data; X. Liu and A. Gennerich wrote the manuscript. A. Gennerich and W. Qiu secured funding.

## Methods

### Plasmids

The pET-17b plasmid containing the codon-optimized GFP-KlpA construct was generated as described previously^17^. The mCherry-KlpA construct was derived by replacing the N-terminal GFP of GFP-KlpA with the sequence of mCherry. The full-length HEST and the tail-truncated HSET (HSETΔTail) constructs were prepared using the HSET cDNA corresponding to Gen Bank Accession Number BC121041. The full-length HSET construct (amino acids 1-673) and the HSETΔTail construct (amino acids 139-673) were fused with the N-terminal ZZ-Tev tag for purification as well as the GFP sequence for fluorescence microscopy and for binding to anti-GFP antibody-coated trapping beads. These constructs were integrated into the pFastbac vector using Gibson Assembly (NEB, #E5510) for insect cell expression.

### Protein expression and purification from E. *coli*

GFP-KlpA and mCherry-KlpA were expressed in E. *coil* cells as described previously^48,49^. In brief, plasmids were transformed into BL21-CodonPlus (DE3)-RIPL competent cells (Agilent Technologies, #230280) with heat shock at 42 ℃ for 45 seconds followed by incubation at 37 ℃ for 1 hour in SOC medium. The cells were then plated onto LB agar plate containing 50 μg/mL Ampicillin and 30 μg/mL chloramphenicol. Subsequently, a single colony was carefully selected and inoculated in 3 mL of LB Broth for optimal growth. The 3-mL culture was incubated at 37°C overnight with continuous shaking. It was then used to inoculate 400 mL of LB Broth supplemented with 50 μg/mL Ampicillin and 30 μg/mL chloramphenicol. The 400-mL culture was incubated at 37°C with vigorous shaking for 4-5 hours and subsequently cooled on ice for 1 hour. To induce expression, IPTG was added to the culture at a final concentration of 0.1 mM. The culture was further incubated at 18°C overnight while being shaken. The cells were harvested by centrifugation at 3,000 relative centrifugal force for 10 minutes at 4°C. Following the removal of the supernatant, the cell pellet was fully resuspended in 5 mL of B-PER complete bacterial protein extraction reagent (Thermo Fisher Scientific, #89821) containing 2 mM MgCl_2_, 1 mM EGTA, 1 mM DTT, 0.1 mM ATP, 2 mM PMSF, and 10% glycerol. The fully resuspended cells were flash-frozen and stored at −80°C for further use.

The frozen cells were thawed at 37°C to initiate protein purification. Following thawing, the solution was nutated at room temperature for 20 minutes to promote thorough mixing. The cells were then gently lysed by douncing on ice with 10 strokes. The resulting cell lysate was clarified by centrifugation at 80,000 rpm for 10 minutes at 4°C using a Beckman Coulter tabletop centrifuge unit. The lysate supernatant was mixed with 200 µL of Ni-nitrilotriacetic acid resin (Roche Complete His-Tag purification resin, #5893682001; MilliporeSigma) and incubated at 4°C with gentle nutation for 1 hour. After incubation with the Ni-nitrilotriacetic acid resin, the resin was washed with wash buffer (50 mM Hepes, 300 mM KCl, 2 mM MgCl_2_, 1 mM EGTA, 1 mM DTT, 1 mM PMSF, 0.1 mM ATP, 0.1% Triton X-100, 10% glycerol, pH 7.2). Then, the protein was eluted using elution buffer (wash buffer with 250 mM imidazole). The eluate was flash-frozen and stored at −80°C for further use.

### Protein expression and purification from insect cell

HSET and HSETΔTail were expressed in Sf9 cells as described previously^48^. Briefly, the pFastBac plasmid containing either HSET or HSETΔTail was transformed into DH10Bac-competent cells (Gibco, #10361012) using a heat-shock method. The transformation was performed by subjecting the cells to heat shock at 42°C for 45 seconds, followed by incubation at 37°C for 4 hours in S.O.C. medium (Gibco, #15544034). Subsequently, the transformed cells were plated onto LB agar plates supplemented with kanamycin (50 μg/mL), gentamicin (7 μg/mL), tetracycline (10 μg/mL), BluoGal (100 μg/mL), and isopropyl-β-D-thiogalactoside (IPTG, 40 μg/mL). After a 36-hour incubation period, positive clones were identified based on the blue/white color screen. Subsequently, a single white colony was carefully selected and inoculated in 3 mL LB broth supplemented with kanamycin (50 μg/mL), gentamicin (7 μg/mL), tetracycline (10 μg/mL) followed by incubation at 37°C overnight. Bacmid DNA was extracted from overnight culture using an isopropanol precipitation method with QIAGEN buffer (QIAGEN, #27104) as described previously^48^. For the generation of baculovirus intended for Sf9 insect cell transfection, 2 mL of Sf9 cells at a density of 0.5 × 10^6^ cells per mL were seeded into six-well plates (Corning, #3516). Subsequently, transfection was carried out by adding 2 μg of freshly prepared bacmid DNA and 6 μL of FuGene HD transfection reagent (Promega, E2311), following the instructions provided by the manufacturer. The cells were incubated for 4 days, and the supernatant containing V0 virus was collected. To generate V1 virus, 0.5 mL of V0 virus was used to transfect 50 mL of Sf9 cells at 1.5 × 10^6^ cells per mL. The supernatant containing V1 virus was collected by centrifugation at 200 × g for 5 minutes at 4°C after 3 days. The V1 virus was stored at 4°C in the dark until use. To initiate protein expression, 5 mL of the V1 virus was utilized to transfect 500 mL of Sf9 cells at a density of 2 × 10^6^ cells per mL. Following a 60 hours-incubation period, the cells were harvested by centrifugation at 3000 × g for 10 minutes at 4°C. The resulting cell pellet was resuspended in 15 mL of ice-cold PBS and subjected to another round of centrifugation. After removing the supernatant, the cell pellet was flash-frozen in liquid nitrogen and stored at −80°C for subsequent use.

HSET and HSETΔTail were purified from frozen Sf9 pellets as described previously^48^. The frozen pellets from a 500 mL insect cell culture were thawed on ice and resuspended in lysis buffer (50 mM HEPES pH 7.4, 2 mM MgCl_2_, 1 mM EGTA, 100 mM NaCl, 1 mM DTT, 0.1 mM ATP, 10% [v/v] glycerol, 2 mM PMSF) supplemented with one protease inhibitor cocktail tablet (cOmplete-EDTA free, Roche, #11836170001) to a final volume of 50 mL. The cells were lysed using a Dounce homogenizer with 20 strokes, and the resulting lysate was cleared by centrifugation at 279,288 × g for 10 minutes at 4°C using a tabletop centrifuge unit from Beckman Coulter. The clarified supernatant was incubated with 3 mL of IgG Sepharose 6 Fast Flow beads (Cytiva, #17096901) for 4 hours with rotation. Following the incubation period, the protein-bound IgG beads were transferred to a gravity flow column. The beads were then washed sequentially with 100 mL of lysis buffer and 100 mL of TEV buffer (50 mM Tris–HCl pH 8.0, 250 mM KAc, 2 mM MgCl_2_, 1 mM EGTA, 1 mM DTT, 0.1 mM Mg-ATP and 10% [v/v] glycerol). Subsequently, the beads were resuspended in TEV buffer with a final volume of 5 mL. Then, 100 μL of TEV protease (New England Biolabs, #P8112S) was added, and the mixture was incubated at 4°C on a roller overnight. After TEV cleavage, the beads were separated by centrifugation, and the protein of interest was concentrated to 1 mL using a 50 kDa molecular weight cut-off (MWCO) centrifugal filter (Millipore, #UFC505024). Finally, the concentrated protein was flash-frozen in liquid nitrogen for storage.

### MT-binding and -release assay

MT-binding and -release purification was performed to further purify the motor proteins and to remove any inactive motors and aggregates. 50 µL of purified protein was buffer exchanged into low-salt buffer (30 mM Hepes, 50 mM KCl, 2 mM MgCl_2_, 1 mM EGTA, 1 mM DTT, and 0.1 mM AMPPNP) using a 0.5-mL Zeba spin desalting column (7-kD molecular weight cutoff, Thermo Fisher Scientific, #89882). AMP-PNP and Taxol were added to the flow-through to final concentrations of 1 mM and 10 µM, respectively. After adding 5 µL of 5 mg/mL Taxol-stabilized MTs to the mixture, the solution was incubated at room temperature for 5 minutes to allow motors to bind to the MTs. The mixture was centrifuged through a 100-µL glycerol cushion (80 mM Pipes, 2 mM MgCl_2_, 1 mM EGTA, 1 mM DTT, 10 µM Taxol, and 60% glycerol) at 40,000 rpm for 10 minutes at room temperature. Subsequently, the supernatant was carefully removed, and the pellet was resuspended in 50 µL of high-salt buffer (30 mM Hepes, 300 mM KCl, 2 mM MgCl_2_, 1 mM EGTA, 1 mM DTT, 10 µM Taxol, 3 mM ATP, and 10% glycerol). The MTs were separated by centrifugation at 40,000 rpm for 5 minutes at room temperature. The resulting supernatant, known as the MT-release (MT-R) fraction, was collected and divided into aliquots. The aliquots were then flash-frozen in liquid nitrogen and stored at −80°C for further use.

### Optical tweezers assay

Optical tweezers-based force measurements were performed with a LUMICKS’ C-Trap combined with TIRF. The anti-GFP antibody coated polystyrene trapping beads, microfluidic slide chamber, and MTs were prepared as described previously^37^. Briefly, Rhodamine X-labeled, biotinylated MTs were immobilized onto glass coverslip via the α-casein-biotin and streptavidin. GFP-KlpA, mCherry-KlpA, GFP-HSET or GFP-HSETΔTail were diluted to the appropriate concentrations using motility buffer (30 mM Hepes, 50 mMKAc, 2 mM MgCl_2_, 1 mM EGTA, 1 mM DTT, 10 µM Taxol, 2 mM ATP, 50 mM glucose, gloxy, 0.75 mg/mL α-casein, pH 7.2) and incubated with anti-GFP antibody-coated, ∼1 μm-diameter beads (990 nm, carboxyl-modified polystyrene microspheres, Polysciences, #08226-15) for 10 minutes. The mixture was then supplemented with 1 mM ATP and flown into the slide chamber. Trapping assays were performed at 25°C with trap stiffnesses of 0.04–0.3 pN/nm.

### MT-gliding assay

The flow chamber was assembled as described previously^50^. To immobilize motors on the coverslip surface, 12 μL of a 0.4 mg/mL rabbit monoclonal anti-GFP antibody solution was introduced into the slide chamber and incubated for 10 minutes. The chamber was then washed twice with 20 μL of blocking buffer, which consisted of 80 mM PIPES, 2 mM MgCl_2_, 1 mM EGTA, 1% Pluronic F-127, and 1 mg/mL α-casein. The blocking buffer was incubated for an additional 10 minutes to block the glass surface. After the blocking step, the slide chamber was washed twice with 20 μL motility buffer. Subsequently, 10 μL of the motor solution (appropriately diluted in motility buffer) was introduced into the slide chamber and incubated for 2 minutes. The flow chamber was then washed twice with 20 μL motility buffer to remove unbound motors. A final 20 μL motility buffer containing fluorescently labeled MTs and supplied with 1 mM ATP, 20 μM Taxol and 1 μL of oxygen-scavenger system were introduced into the chamber. MTs were visualized with a custom-built TIRF microscope and imaged with an acquisition time of 200 ms for a total 200 frames per movie. The velocity was analyzed using Fiji and the statistical analysis and data visualization were performed using Prism.

## References

1. Walczak, C.E. & Heald, R. Mechanisms of mitotic spindle assembly and function. Int Rev Cytol 265, 111–58 (2008).

2. Peterman, E.J. & Scholey, J.M. Mitotic microtubule crosslinkers: insights from mechanistic studies. Curr Biol 19, R1089–94 (2009).

3. Cross, R.A. & McAinsh, A. Prime movers: the mechanochemistry of mitotic kinesins. Nat Rev Mol Cell Biol 15, 257–71 (2014).

4. Pandey, H., Popov, M., Goldstein-Levitin, A. & Gheber, L. Mechanisms by Which Kinesin-5 Motors Perform Their Multiple Intracellular Functions. Int J Mol Sci 22(2021).

5. Mann, B.J. & Wadsworth, P. Kinesin-5 Regulation and Function in Mitosis. Trends Cell Biol 29, 66–79 (2019).

6. Impastato, A.C. et al. Optical Control of Mitosis with a Photoswitchable Eg5 Inhibitor. Angew Chem Int Ed Engl 61, e202115846 (2022).

7. Valentine, M.T., Fordyce, P.M., Krzysiak, T.C., Gilbert, S.P. & Block, S.M. Individual dimers of the mitotic kinesin motor Eg5 step processively and support substantial loads in vitro. in Nat Cell Biol Vol. 8 470–476 (2006).

8. She, Z.Y. & Yang, W.X. Molecular mechanisms of kinesin-14 motors in spindle assembly and chromosome segregation. J Cell Sci 130, 2097–2110 (2017).

9. Yamada, M., Tanaka-Takiguchi, Y., Hayashi, M., Nishina, M. & Goshima, G. Multiple kinesin-14 family members drive microtubule minus end-directed transport in plant cells. J Cell Biol 216, 1705–1714 (2017).

10. Hepperla, A.J. et al. Minus-end-directed Kinesin-14 motors align antiparallel microtubules to control metaphase spindle length. Dev Cell 31, 61–72 (2014).

11. Cai, S., Weaver, L.N., Ems-McClung, S.C. & Walczak, C.E. Kinesin-14 family proteins HSET/XCTK2 control spindle length by cross-linking and sliding microtubules. Mol Biol Cell 20, 1348–59 (2009).

12. Olmsted, Z.T., Colliver, A.G., Riehlman, T.D. & Paluh, J.L. Kinesin-14 and kinesin-5 antagonistically regulate microtubule nucleation by gamma-TuRC in yeast and human cells. Nat Commun 5, 5339 (2014).

13. Kim, N. & Song, K. KIFC1 is essential for bipolar spindle formation and genomic stability in the primary human fibroblast IMR-90 cell. Cell Struct Funct 38, 21–30 (2013).

14. Zhu, C. et al. Functional analysis of human microtubule-based motor proteins, the kinesins and dyneins, in mitosis/cytokinesis using RNA interference. Mol Biol Cell 16, 3187–99 (2005).

15. Hatsumi, M. & Endow, S.A. Mutants of the microtubule motor protein, nonclaret disjunctional, affect spindle structure and chromosome movement in meiosis and mitosis. J Cell Sci 101 **(Pt** **3**), 547–59 (1992).

16. Acilan, C. & Saunders, W.S. A tale of too many centrosomes. Cell 134, 572–5 (2008).

17. Popchock, A.R. et al. The mitotic kinesin-14 KlpA contains a context-dependent directionality switch. Nat Commun 8, 13999 (2017).

18. Endres, N.F., Yoshioka, C., Milligan, R.A. & Vale, R.D. A lever-arm rotation drives motility of the minus-end-directed kinesin Ncd. in Nature Vol. 439 875–878 (2006).

19. Furuta, K. et al. Measuring collective transport by defined numbers of processive and nonprocessive kinesin motors. Proc Natl Acad Sci U S A 110, 501–6 (2013).

20. Braun, M. et al. Changes in microtubule overlap length regulate kinesin-14-driven microtubule sliding. Nat Chem Biol 13, 1245–1252 (2017).

21. Norris, S.R. et al. Microtubule minus-end aster organization is driven by processive HSET-tubulin clusters. Nat Commun 9, 2659 (2018).

22. Wendt, T.G. et al. Microscopic evidence for a minus-end-directed power stroke in the kinesin motor ncd. EMBO J 21, 5969–78 (2002).

23. deCastro, M.J., Fondecave, R.M., Clarke, L.A., Schmidt, C.F. & Stewart, R.J. Working strokes by single molecules of the kinesin-related microtubule motor ncd. in Nat Cell Biol Vol. 2 724–729 (2000).

24. Endow, S.A. & Higuchi, H. A mutant of the motor protein kinesin that moves in both directions on microtubules. in Nature Vol. 406 913–916 (2000).

25. Reinemann, D.N., Norris, S.R., Ohi, R. & Lang, M.J. Processive Kinesin-14 HSET Exhibits Directional Flexibility Depending on Motor Traffic. Curr Biol 28, 2356–2362 e5 (2018).

26. Wang, P. et al. The Central Stalk Determines the Motility of Mitotic Kinesin-14 Homodimers. Curr Biol 28, 2302–2308 e3 (2018).

27. Yun, M. et al. Rotation of the stalk/neck and one head in a new crystal structure of the kinesin motor protein, Ncd. EMBO J 22, 5382–9 (2003).

28. Nitzsche, B. et al. Working stroke of the kinesin-14, ncd, comprises two substeps of different direction. Proc Natl Acad Sci U S A 113, E6582–E6589 (2016).

29. Kozielski, F., De Bonis, S., Burmeister, W.P., Cohen-Addad, C. & Wade, R.H. The crystal structure of the minus-end-directed microtubule motor protein ncd reveals variable dimer conformations. Structure 7, 1407–16 (1999).

30. Sablin, E.P. et al. Direction determination in the minus-end-directed kinesin motor ncd. Nature 395, 813–6 (1998).

31. Kaneko, T. et al. Different motilities of microtubules driven by kinesin-1 and kinesin-14 motors patterned on nanopillars. Sci Adv 6, eaax7413 (2020).

32. Furuta, K.a.y., Edamatsu, M., Maeda, Y. & Toyoshima, Y.Y. Diffusion and directed movement: in vitro motile properties of fission yeast kinesin-14 Pkl1. in J Biol Chem Vol. 283 36465–36473 (2008).

33. Allersma, M.W., Gittes, F., deCastro, M.J., Stewart, R.J. & Schmidt, C.F. Two-dimensional tracking of ncd motility by back focal plane interferometry. Biophys J 74, 1074–1085 (1998).

34. Butterfield, A.E., Stewart, R.J., Schmidt, C.F. & Skliar, M. Bidirectional power stroke by ncd kinesin. Biophys J 99, 3905–15 (2010).

35. Brenner, S., Berger, F., Rao, L., Nicholas, M.P. & Gennerich, A. Force production of human cytoplasmic dynein is limited by its processivity. Sci Adv 6, eaaz4295 (2020).

36. Elshenawy, M.M. et al. Cargo adaptors regulate stepping and force generation of mammalian dynein-dynactin. Nat Chem Biol 15, 1093–1101 (2019).

37. Budaitis, B.G. et al. Pathogenic mutations in the kinesin-3 motor KIF1A diminish force generation and movement through allosteric mechanisms. J Cell Biol 220(2021).

38. Svoboda, K. & Block, S.M. Force and velocity measured for single kinesin molecules. Cell 77, 773–784 (1994).

39. Lam, A.J. et al. A highly conserved 310 helix within the kinesin motor domain is critical for kinesin function and human health. Sci Adv 7(2021).

40. Plotnikov, S.V., Millard, A.C., Campagnola, P.J. & Mohler, W.A. Characterization of the myosin-based source for second-harmonic generation from muscle sarcomeres. Biophys J 90, 693–703 (2006).

41. Llewellyn, M.E., Barretto, R.P., Delp, S.L. & Schnitzer, M.J. Minimally invasive high-speed imaging of sarcomere contractile dynamics in mice and humans. Nature 454, 784–8 (2008).

42. Spudich, J.A. Hypertrophic and dilated cardiomyopathy: four decades of basic research on muscle lead to potential therapeutic approaches to these devastating genetic diseases. Biophys J 106, 1236–49 (2014).

43. Matusovsky, O.S., Mansson, A. & Rassier, D.E. Cooperativity of myosin II motors in the non-regulated and regulated thin filaments investigated with high-speed AFM. J Gen Physiol 155(2023).

44. Siddiqui, N. et al. Force generation of KIF1C is impaired by pathogenic mutations. Curr Biol 32, 3862–3870 e6 (2022).

45. Shimamoto, Y., Forth, S. & Kapoor, T.M. Measuring Pushing and Braking Forces Generated by Ensembles of Kinesin-5 Crosslinking Two Microtubules. Dev Cell 34, 669–81 (2015).

46. Svoboda, K., Schmidt, C.F., Schnapp, B.J. & Block, S.M. Direct observation of kinesin stepping by optical trapping interferometry. Nature 365, 721–727 (1993).

47. Yildiz, A., Tomishige, M., Vale, R.D. & Selvin, P.R. Kinesin walks hand-over-hand. Science 303, 676–8 (2004).

48. Fu, X. et al. Doublecortin and JIP3 are neural-specific counteracting regulators of dynein-mediated retrograde trafficking. Elife 11(2022).

49. Pant, D.C. et al. ALS-linked KIF5A DeltaExon27 mutant causes neuronal toxicity through gain-of-function. EMBO Rep 23, e54234 (2022).

50. Liu, X., Rao, L. & Gennerich, A. The regulatory function of the AAA4 ATPase domain of cytoplasmic dynein. Nat Commun 11, 5952 (2020).

