## Supplemental Figures for "Kinesin-14 HSET and KlpA are non-processive microtubule motors with load-dependent power strokes"

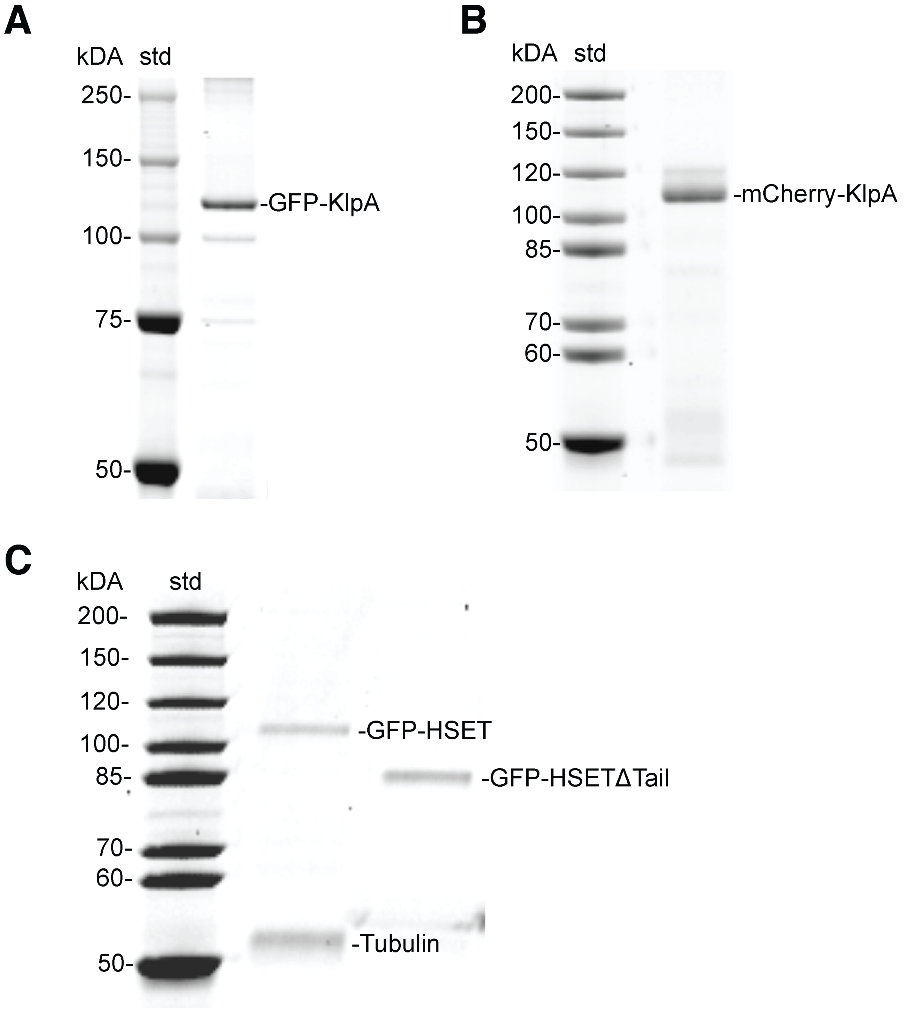


**Suppl. Fig. 1 SDS-PAGE gel.** (A) SDS-PAGE of molecular mass marker (left) and *E. Coli*-expressed GFP-KlpA (right); (B) SDS-PAGE of molecular mass marker (left) and *E. Coli*-expressed mCherry-KlpA (right); (C) SDS-PAGE of molecular mass marker (left), insect cell-expressed and MT-binding and -release purified GFP-HSET (middle), and GFP-HSET-ΔTail (right).

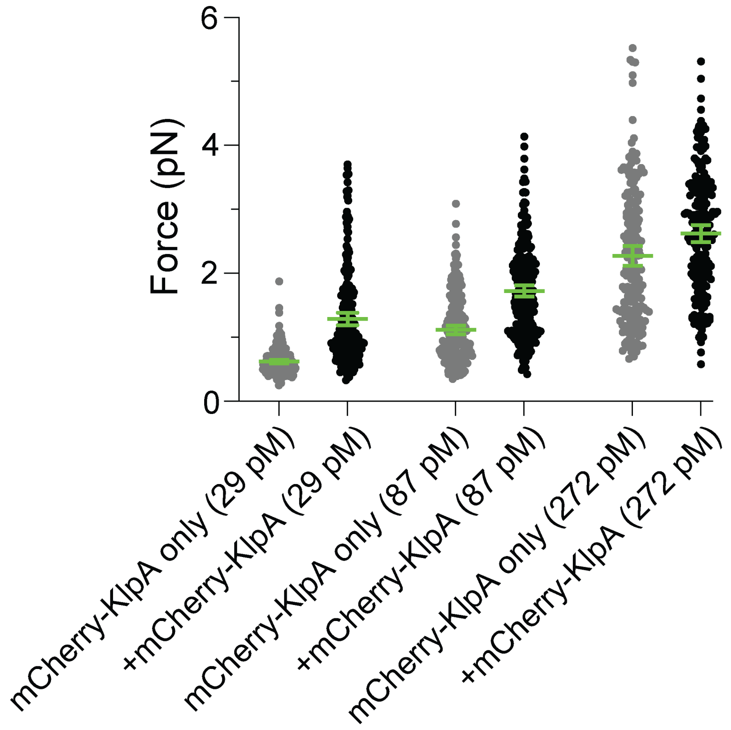


**Suppl. Fig. 2 KlpA force generation as a function of increasing concentrations of mCherry-KlpA in the absence and presence of pre-bound GFP-KlpA.** Control experiment comparing the force differences of anti-GFP antibody-coated beads without (grey dots) and with pre-bound GFP-KlpA (black dots) in the presence of increasing concentrations of mCherry-KlpA at $k$=0.05 pN/nm. Green bars represent the mean values with 95% CIs. 29 pM mCherry-KlpA alone: 0.62 ± 0.01 pN pN ($n$=232); GFP-KlpA at the single-molecule level (14 pM) prebound to beads with added 29 pM mCherry-KlpA: 1.29 ± 0.05 pN ($n$=232); 87 pM mCherry-KlpA alone: 1.12 ± 0.03 pN ($n$=244); 14 pM GFP-KlpA plus 87 pM mCherry-KlpA: 1.73 ± 0.05 pN ($n$=259); 272 pM mCherry-KlpA alone: 2.27 ± 0.08 pN ($n$=188); 14 pM GFP-KlpA plus 272 pM mCherry-KlpA: 2.62 ± 0.07 pN ($n$=203).

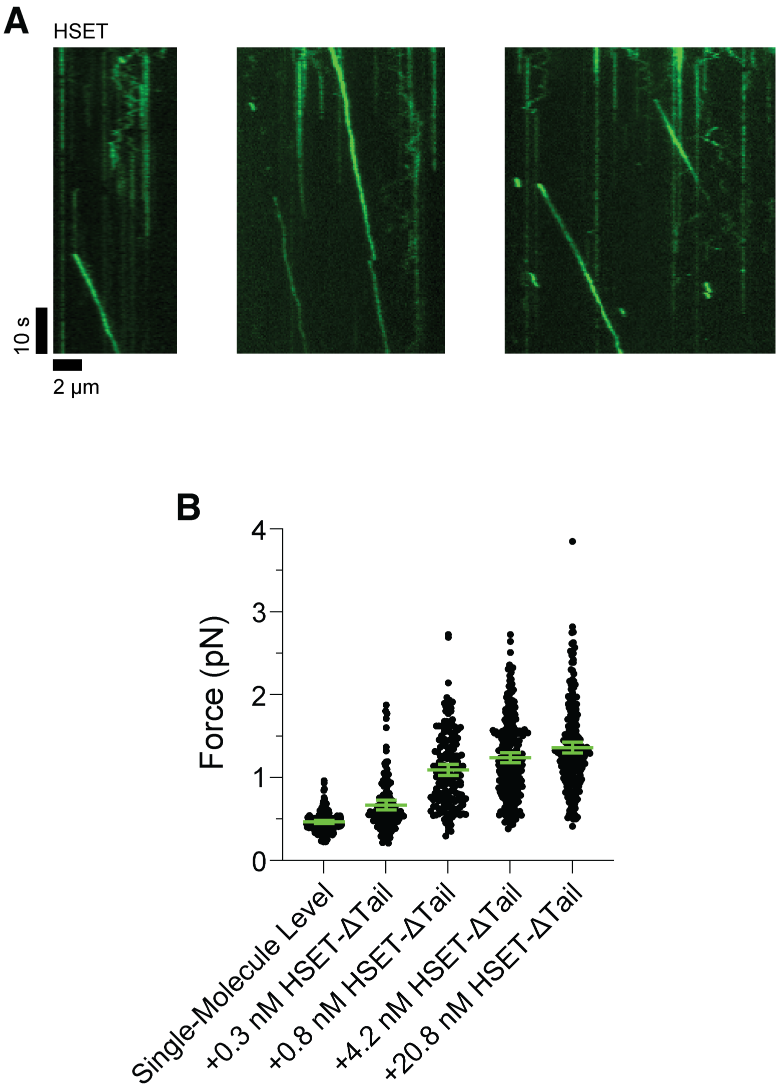


**Suppl. Fig. 3 Processive motion of oligomerized HSET and force generation of GFP-HSET-ΔTail at increasing motor concentrations.** (A) Examples of MT minus-end-directed processive motion of HSET particles formed at 1 nM motor concentration. These particles consist of multiple HSET motors working together. (B) Forces generated by GFP-HSET-ΔTail with increasing motor concentrations at $k$=0.1 pN/nm. Green bars represent the mean values with 95% CIs. GFP-HSET-ΔTail at single-molecule level (30 pM): 0.46 ± 0.01 pN ($n$=197); +0.33 nM GFP-HSET-ΔTail: 0.67 ± 0.03 pN ($n$=124); +0.83 nM GFP-HSET-ΔTail: 1.09 ± 0.03 pN ($n$=170); +4.15 nM GFP-HSET-ΔTail: 1.24 ± 0.03 pN ($n$=252); +20.8 mM GFP-HSET-ΔTail: 1.36 ± 0.03 pN ($n$=236).

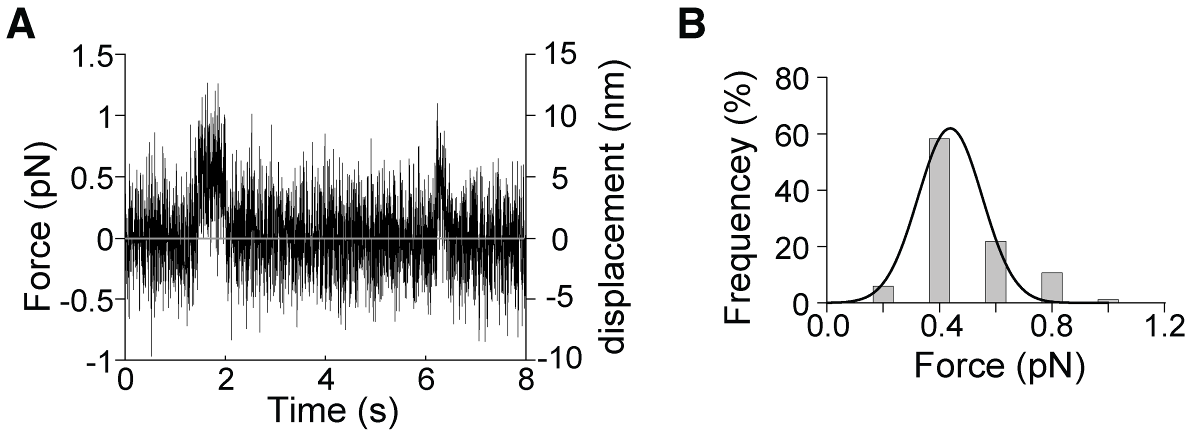


**Suppl. Fig. 4 Force generation of single *E. coli*-expressed GFP-HSET molecules.** (A) Representative force-versus-time trace depicting bead movement driven by a single *E. coli*-expressed GFP-HSET molecule at 1 mM ATP and $k$=0.1 pN/nm. (B) Histogram of forces generated by single GFP-HSET molecules (0.44 ± 0.014 pN, mean ± SEM from Gaussian fit, $n$=63).
